# HPLC analysis of *Withania somnifera* (Linn.) Dunal reveals two chemotypes in Indian germplasm and seasonal and genetic variability in Withaferin A -Content

**DOI:** 10.1101/2020.07.31.231316

**Authors:** Nutan Kaushik

## Abstract

HPLC methods for profiling and estimation of withanolide in *Withania* have been developed to study seasonal, geographical and genetic variation in Withaferin-A content and identify chemotypes. Highest withanolide content was observed during the vegetative phase and slight decrease was observed during reproductive phase. However, the content drastically reduced at the maturity. No significant effect of location was observed on the withanolide content when same genotypes were grown at two diverse locations. Withaferin-A content in Indian population of *Withania somnifera* varied from 0.22 to 3.38 % with a mean value of 1.099 in the leaves during vegetative stage.Whereas during maturity the content ranged from 0.0056 to 1.99 % with a mean value of 0.5942 %. In the roots, Withaferin-A content ranged from 0.0028 to 0.75 % with a mean value of 0.1242 during vegetative phase and 0.0008 to 0.1489 with a mean value of 0.00213 % at maturity of the plant. The population segregated into two chemotypes identified as chemotype-I and as an intermediate of Chemotype–I and Chemotype-II.

---

*Withania somnifera* (Linn.) Dunal commonly known as “Ashwagandha” belongs to the family Solanaceae. Around 1250 species of family Solanaceae have been widely distributed in the warmer parts of whole world. Twenty three species of genus *Withania* are reported among which *Withania somnifera* has high medicinal value. It is an erect, evergreen and perennial shrub that is widely used in traditional Indian medicine for a variety of diseases having adaptogenic, tonic, and analgesic, antipyretic, anti-inflammatory and abortifacient properties and is one of the most extensively used plants in Ayurvedic and Unani medicine^1^. Both leaves and roots of the plant are used as a source for drug and steroidal lactones.

*W. somnifera* has a fairly wide geographic distribution. Besides Indian subcontinent, the wild growth of the species has also been reported from Israel, Italy, Pakistan, Afghanistan, Palestine, Egypt, Jordan, Morocco, Spain, Canary Island, Eastern Africa, Congo, Madagascar and South Africa, representing extensive variations of soil, rainfall, temperature and altitude. In Italy, *W. somnifera* is only present in Sicily and Sardinia, where it is a highly endangered species^2^.

Three chemotypes (I, II, III) are reported in *W. somnifera* differing in their total leaf content of withanolides with various substitution patterns^3,4^.These substitution are characteristic of each chemotype and seem to be genetic in character. The main constituents of chemotype I and chemotype II is Withaferin-A and withanolide D respectively^4^. In chemotype III two groups of compounds have been characterized, one with compounds possessing a normal stereochemistry at C_17_ (i.e. β-oriented side chain) e.g. Withanolide-G and J, and other with *α*-side chain e.g. withanolide-E and F^5^.

Extreme degree of variability in the morphological characteristics and growth habits of *W. somnifera* plants is found in different parts of India^6^. This plant is also cultivated by the farmers. Considering the wide genetic diversity available in India, there is a great scope for screening the rich diversity for chemotyping and Withaferin-A content. Present study was undertaken to develop HPLC method for quality control, identify chemotypes in Indian populations of *Withania somnifera*, and to estimate Withaferin-A content yielding population to harness the full potential of the plant species for cultivation.

## Methodology

### Collection of Leaf samples

Leaf samples were collected from Delhi, India to standardize the analysis of withanolides by HPLC. To study the effect of environmental conditions i.e. soil and agro climatic conditions on the synthesis of secondary metabolites, leaves from five *Withania somnifera* accessions growing in Mandsaur (Madhya Pradesh), India were collected from the same accession growing in The Energy and Resources Institute’s (TERIs) field station at Gwal Pahari, Haryana, India. Leaf samples from these accessions were collected in the month of November i.e. during vegetative phase of the plant (60 Days Old Plant, DOP), January i.e. reproductive phase (120 DOP) and March i.e. at maturity (180 DOP) when withania crop was harvested and analyzed by HPLC. Since no effect of soil and agro climatic condition was observed, in next step, to study the variations in different population of *W. somnifera*, the seeds of the collections made during July to May from different parts of India and then were sown in the field during September at TERI’s field station at Gwal Pahari (Haryana). Plants were identified by Dr Ahmedulla of Botanical Survey of India., Noida, India. The details of the collections are provided in Table 1. Leaf and root samples were collected in November during vegetative phase and in March at maturity for HPLC analysis.

**Table 1.**
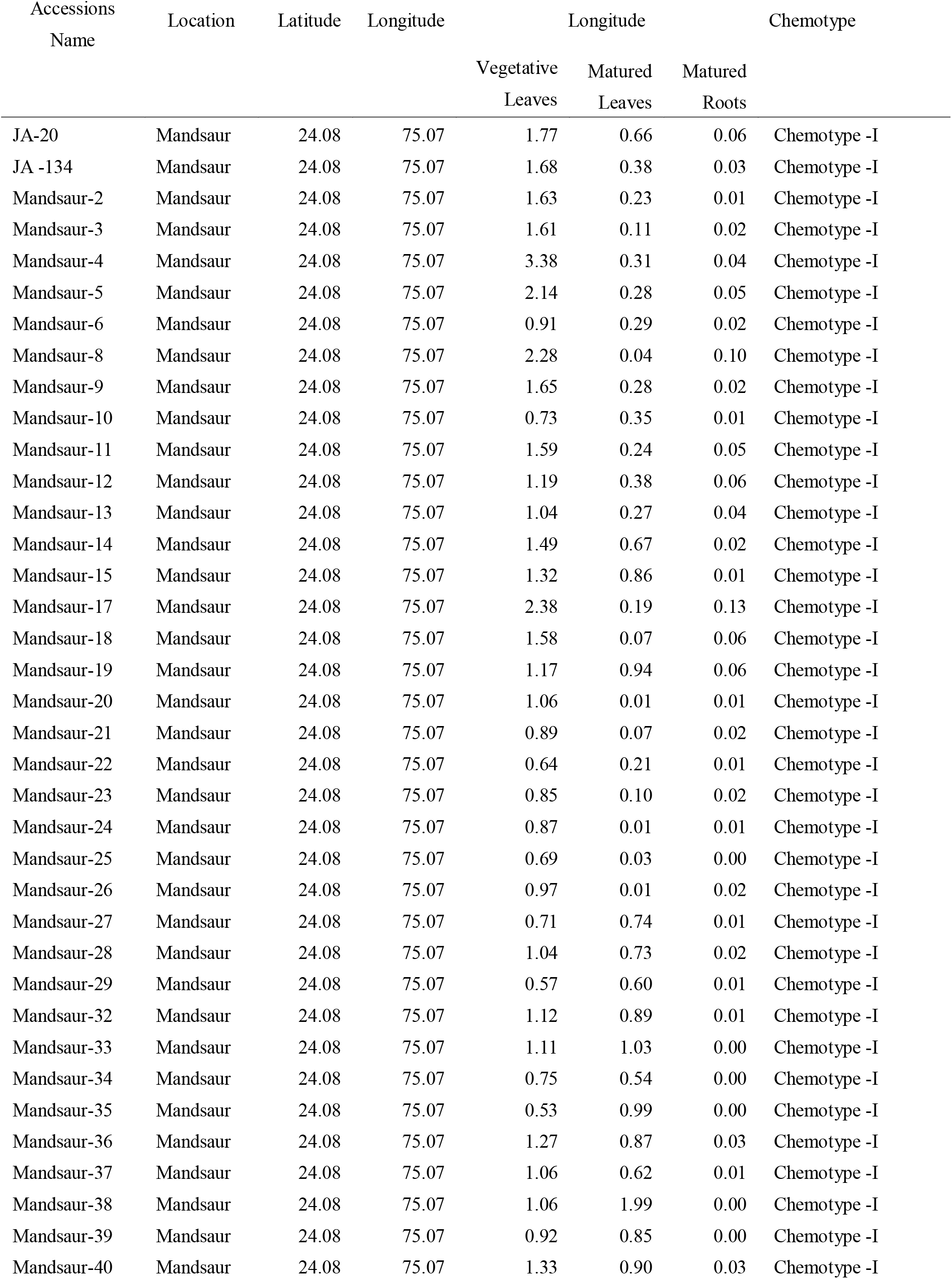
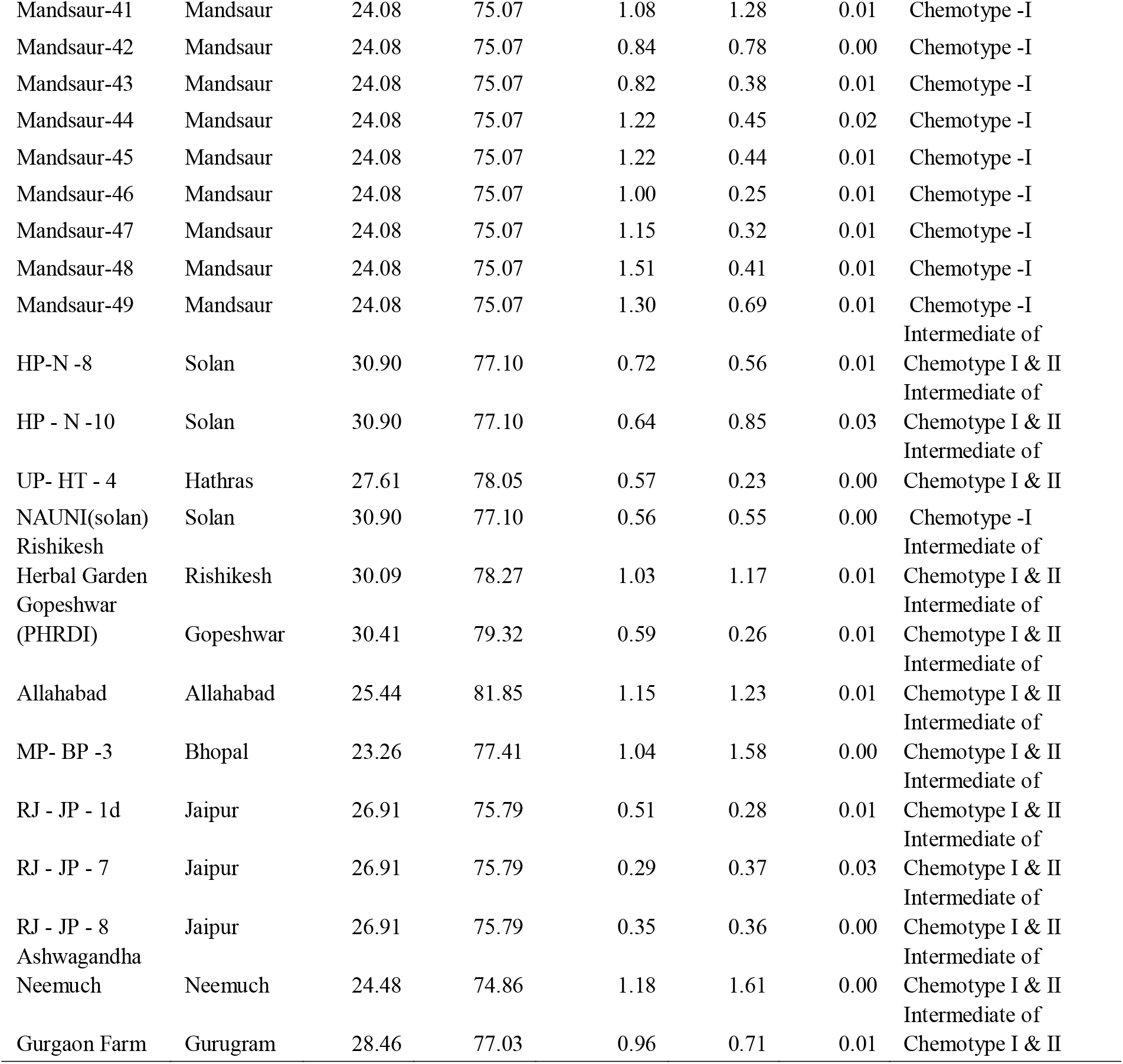
Variability in Withaferin-A content observed in Indianpopulation of *Withania somnifera* collected from different parts of India.

The leaf and root samples were lyophilized by lyophilizer (Virtis, New York, USA) to remove the moisture content. After lyophilization, the leaves and roots were crushed to prepare fine powder in a Kniftech (Germany) grinder.

### Sample preparation for HPLC analysis

Leaf (20 mg) /root powder (50 mg) was weighed accurately in duplicate and transferred into a 15 mL Borosil tube. Distilled ethanol (2 ml) was added to the tube containing the leaf part while 4ml of distilled ethanol was added to the tube containing the root part. The tubes were capped securely and the contents of the tube were heated for 1h on a water bath maintained at 50°C. After 1h, the tubes were removed from the water bath and allowed to cool down to room temperature. The tubes were then centrifuged at 5000 rpm for 5 min and the supernatant was transferred to another tube. The residue left in the tube was re-extracted with distilled ethanol two times (2 mL x 2) by centrifugation. The three supernatants thus obtained were combined together and the volume was made up to 6 mL in the case of leaf samples. However, in the case of root samples the solvent was completely evaporated in a rotary evaporator and then dissolved in 1 mL of HPLC grade methanol (Merck, India). Prepared Samples were filtered through Swinnex polypropylene 25-mm filter holders (Millipore, USA) containing Durapore 0.22 micron filters and kept in 1 mL HPLC vials.

### Preparation of standard solution of Withaferin-A

Pure standard of Withaferin-A (99.99% purity) and withanolide – D (99%) were procured from Chromadex, USA. 1000 ppm standard solution was prepared in HPLC grade methanol. Serial dilutions were performed to obtain standard solutions in the range of 150, 100, 75, 50, 25 and 10 ppm to develop a calibration curve.

### HPLC analysis

Sample and standard (10 μl each) was injected into the Waters HPLC system consisting of M-600 E quaternary gradient pump, 717 autosampler, Novapak Reverse phase C-18 (3.9 × 150 mm) column, and Photo Diode Array (PDA) detector. The separation of the withanolides was obtained on C-18 column using isocratic run of acetonitrile (A) and water (B) in the ratio of 30:70 @ 1 ml/ minute as mobile phase for analysis of Withaferin-A content. For development of the HPLC fingerprints gradient elution was programmed as follows: 0-5 min, 100% B; 5-45 min, 100% B-100% A; 45-50 min, 100% A; 50-60 min, 100% B. The peaks were monitored at 229 nm. The peak identity was confirmed by comparing the retention time of the peaks and also by PDA spectrum. PDA spectrum was also used to confirm the peak purity.

### Quantification of Withaferin-A content

The Withaferin-A content was quantified by using the following formula:

#### 2.5.1. Amount in PPM (μg/g)

Formula:

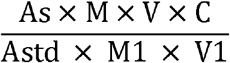

Where,

As= sample area
Astd= standard area
M= μL standard injected
M1= weight of the sample (g)
V= volume of the final extract (mL)
V1= μL sample injected, and
C= concentration in μg per mL of the standard solution

## Results and discussion

### Development of HPLC method and identification of chemotypes

A simple and sensitive HPLC method has been developed for developing HPLC fingerprints of *W. somnifera* leaf and root extracts. The method involves single step extraction in ethanol with no clean up. HPLC fingerprints of leaf and roots of *W. somnifera* were generated by gradient elution (Figure 1).

**Figure 1.**
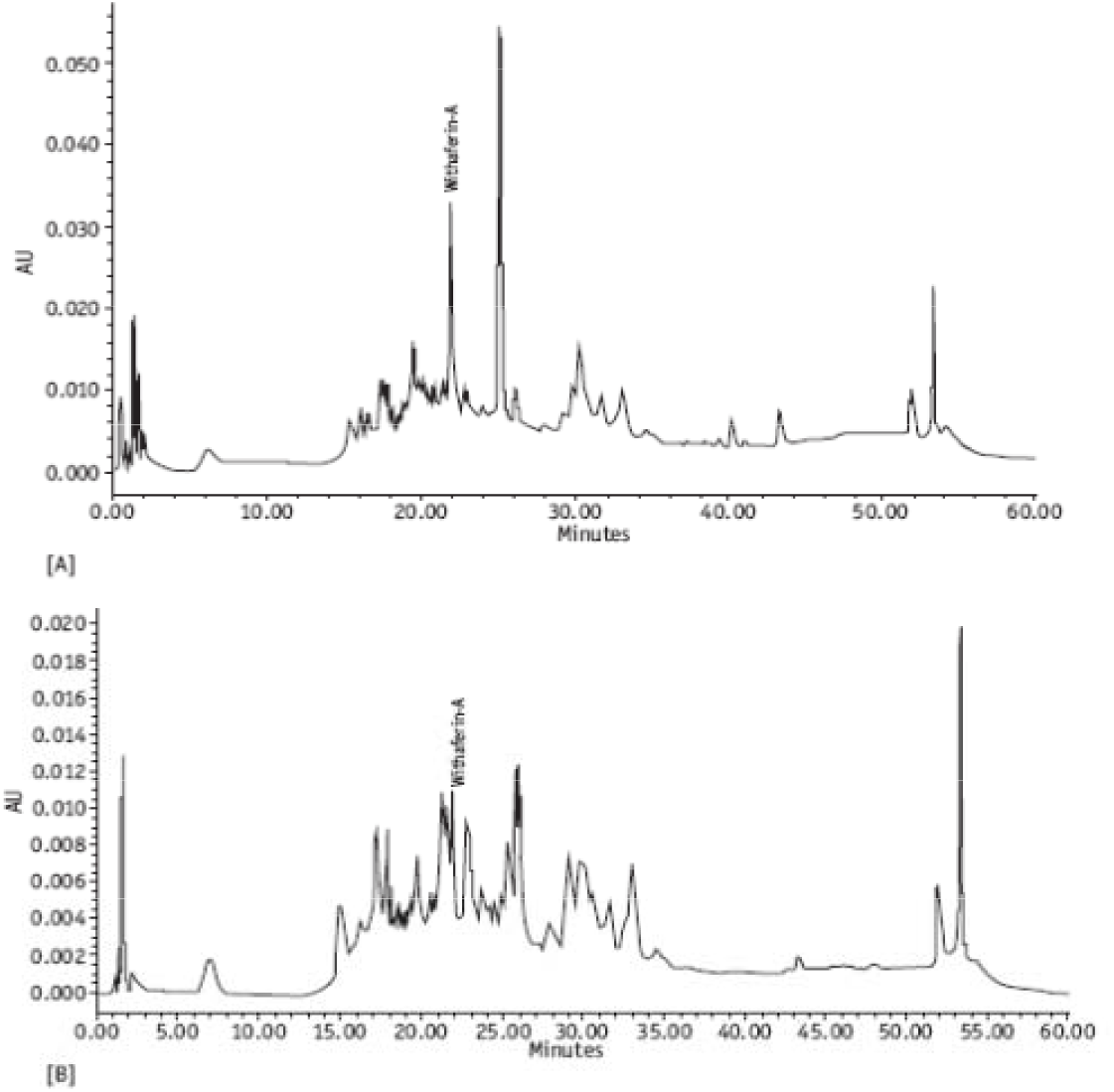
HPLC fingerprint of *Withania somnifera* (A) leaf extract and (B) root extract developed with acetonitrile –water gradient.

HPLC chromatogram obtained with isocratic elution gave reasonably well separation of withanolides for their estimations (Figure 2 and 3). However for for generating the HPLC fingerprint to monitor the qualitygradient elution is recommended. Analysis of the present collections by isocratic method revealed that the cultivated varieties belong to chemotype I having Withaferin-A as main constituent (Figure 2). The germplasm collected from Mandsaur showed Withaferin-A as major constituent. Collection from Yamuna Nagar (Haryana) and Nauni (Himachal Pradesh) also showed the presence of Withaferin-A as main constituent. Collections from Bhopal (Madhya Pradesh), Gopeshwar (Uttranchal), Jaipur (Rajasthan), Delhi, Jodhpur (Rajasthan), Chandigarh (Punjab), and Udaipur (Rajasthan) however showed the dominance of two withanolides viz Withaferin–A and the other identified as withanolide–D in equal proportion thus proved as an intermediate of chemotype-I and chemotype-II (Figure 3).

**Figure 2.**
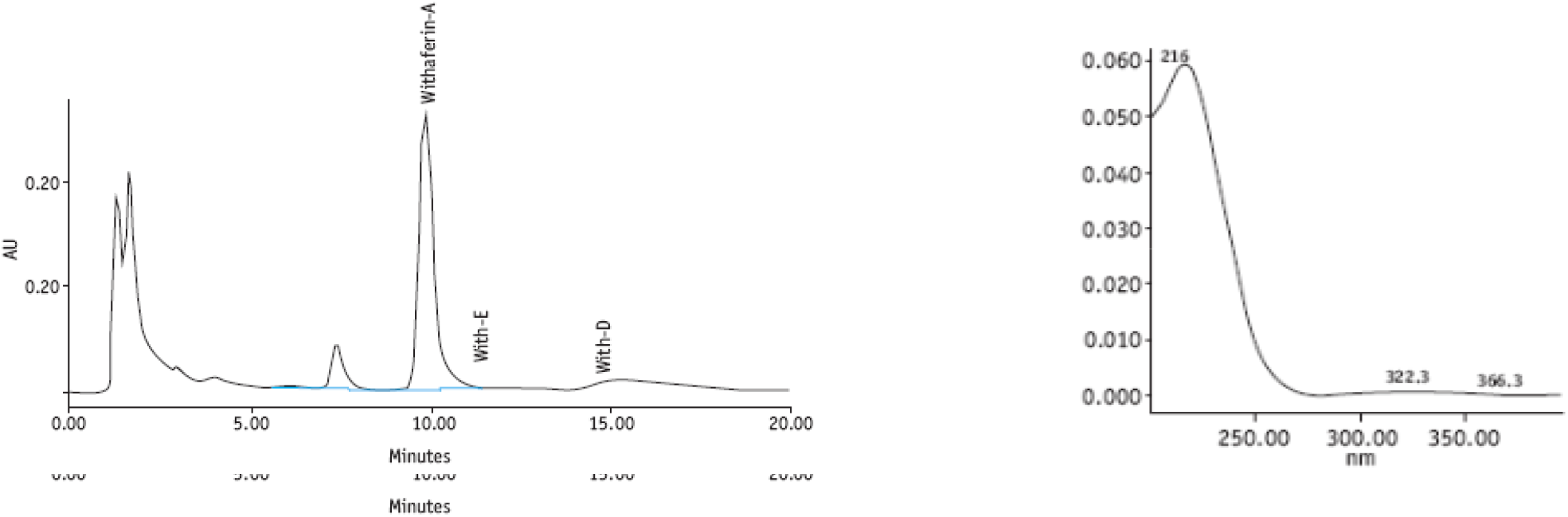
HPLC-PDA chromatogram of *Withania somnifera* leaf extract obtained with isocratic elution of acetonitrile –water (30:70) for estimation of withanolide content showing chemotype I having withaferin-A as main constituent.

**Figure 3.**
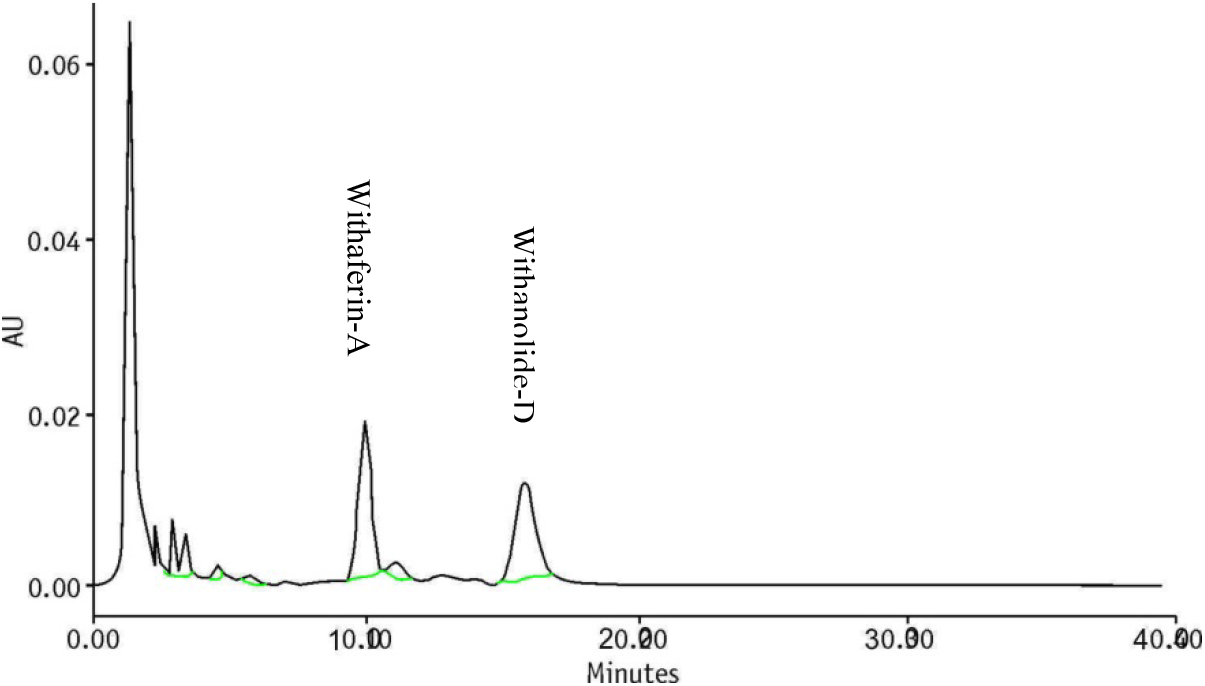
HPLC chromatogram of *Withania somnifera* leaf extract obtained with iscocratic elution intermediate of chemotype I and chemotyp e II having withaferin-A and withanolide –D as main constituents.

### Variations in Withaferin-A Content

Withaferin-A is pharmacologically most important withanolide having anti-inflammatory and immunosuppressive properties. In the present study, variability in Withaferin-A content in the accessions collected from different parts of India was assessed (Table 1). It is extremely important to know the right time for collection and also the effect of environmental conditions on the synthesis of secondary metabolites in order to study the variations in the content. To ensure this, five *Withania somnifera* accessions growing in Mandsaur were selected and the same accessions were also grown in our field station at Gwal Pahari, Haryana. Leaf samples from these accessions were collected in the month of November i.e. during vegetative phase of the plant (60 Days Old Plant, DOP), January i.e. reproductive phase (120 DOP) and March i.e. at maturity (180 DOP) when *Withania* crop was harvested. Withanolide content of these accessions were estimated by HPLC in their leaf. Highest withanolide content was observed in November samples i.e. during the vegetative phase and a slight decrease was observed in January collection i.e. the reproductive phase (Table 2). However, the content drastically reduced in March samples at the maturity stage. On an average, 28.74% decrease in Withaferin-A content was observed during reproductive phase and 81.04% decrease at maturity as compared to the Withaferin-A content during vegetative phase in the leaves. No significant effect of location was observed on the withanolide content. The soil type of Mandsaur is black cotton soil whereas Gwal Pahari has clayey soil. Thus the withanolide content remained unaffected by the environment particularly the soil type.

**Table 2.** Withaferin-A content (% dry weight) of the *Withania somnifera* leaves collected after different days of sowing in different populations.

| leaves collected after different days of sowing in different populations. |  |  |  |  |  |  |
| --- | --- | --- | --- | --- | --- | --- |
| Withaferin A (%) |  |  |  |  |  |  |
| Collection No | Mandsaur |  |  | Delhi |  |  |
|  | 60 DOP | 120 DOP | 150 DOP | 60 DOP | 120 DOP | 150 DOP |
| JA-20 | 1.63 | 1.64 | 0.66 | 1.77 | 1.66 | 0.66 |
| JA-134 | 1.71 | 1.17 | 0.35 | 1.68 | 1.20 | 0.38 |
| Mandsaur-2 | 1.65 | 1.31 | 0.25 | 1.63 | 1.28 | 0.23 |
| Mandsaur-51 | 1.62 | 1.32 | 0.23 | 1.54 | 1.37 | 0.22 |
| Mandsaur-3 | 1.58 | 1.10 | 0.18 | 1.61 | 0.86 | 0.11 |

\*DOP = Days old plant

In view of these observations seeds collected from different accessions of withania during July-May were sown in the field during September at TERI’s field station at Gwal Pahari (Haryana). Some of the seeds did not germinate. Since drastic decrease in Withaferin-A content was observed at the time of harvest, it was decided to study the Withaferin-A content not only in leaves but also in roots. It was done in order to see whether Withaferin-A content was getting translocated from the leaves to the roots at the time of maturity. Therefore, leaf and root samples were collected during vegetative phase in November and at maturity i.e. in May, in order to estimate the variability in terms of withanolide content.

The analysis of these samples revealed the wide variability in terms of Withaferin-A content in the collected accessions. The Withaferin-A content varied from 0.22 to 3.38 % with a mean value of 1.099% in the leaves during vegetative stage while during maturity the content ranged from 0.0056 to 1.99 % with a mean value of 0.5942 % (Table 3). In the roots, Withaferin-A content ranged from 0.0028 to 0.75 % with a mean value of 0.1242 during vegetative phase and 0.0008 to 0.1489 with a mean value of 0.00213 % at maturity of the plant. This indicates that the leaf Withaferin-A is not being translocated from leaf to roots as decrease in Withaferin-A content was observed both in leaves and in roots (Figure 4). Further, the percentage decrease of Withaferin-A content in roots was higher with an average value of 90% than the decrease observed in the leaves. Interestingly, few plants did not show significant decrease in Withaferin-A content in leaves at the time of maturity (Table 4). Table 1 provides the variations observed in Withaferin-A content in different accessions. The highest Withaferin-A content (3.38%) was recorded in the plants collected from Mandsaur and the lowest (0.23%) in the plants collected from Hathras (Uttar Pradesh). The Withaferin content of the popular variety of the withania Jawahar Ashwagandha-20 (JA-20) was estimated as 1.77 %. Frequency histograms are provided in Figure. 5. Scattered plot obtained with this population (Figure 6) clearly demarcated the root and leaf content and further confirmed that Withaferin–A content in leaves was higher as compared to root. This was irrespective of age and expression of Withaferin-A, which was higher during 60 DOP i.e. vegetative phase of the plant.

**Figure 4.**
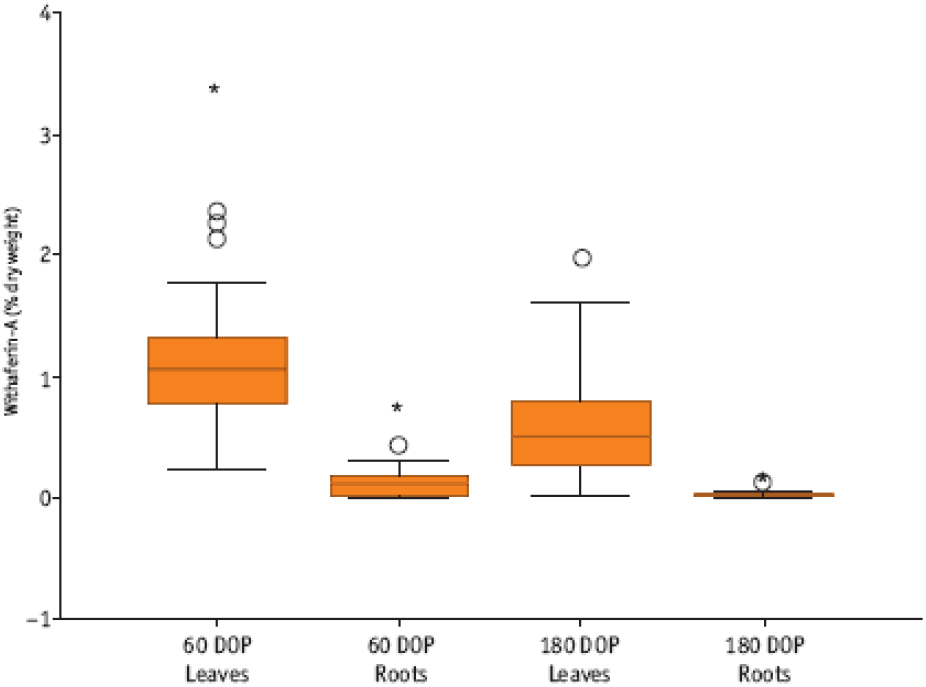
Comparison of means of Withafern-A (% dry weight) in leaves and roots at different physiological stages of the *W. somnifera* plant population.

**Figure 5.**
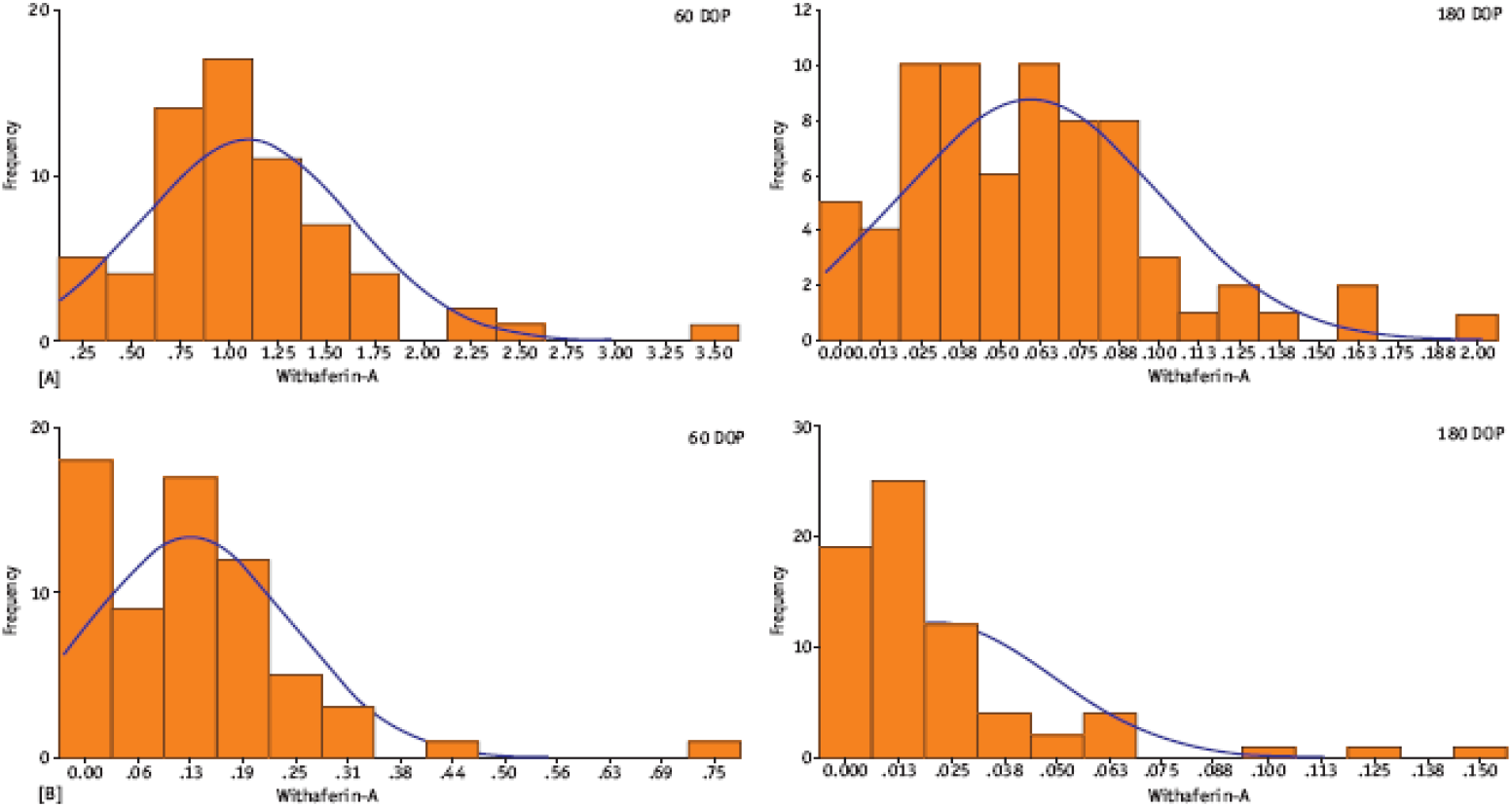
Frequency histogram for withaferin-A content (% dry weight) in leaves and roots at different physiological stages of the plant.

**Figure 6.**
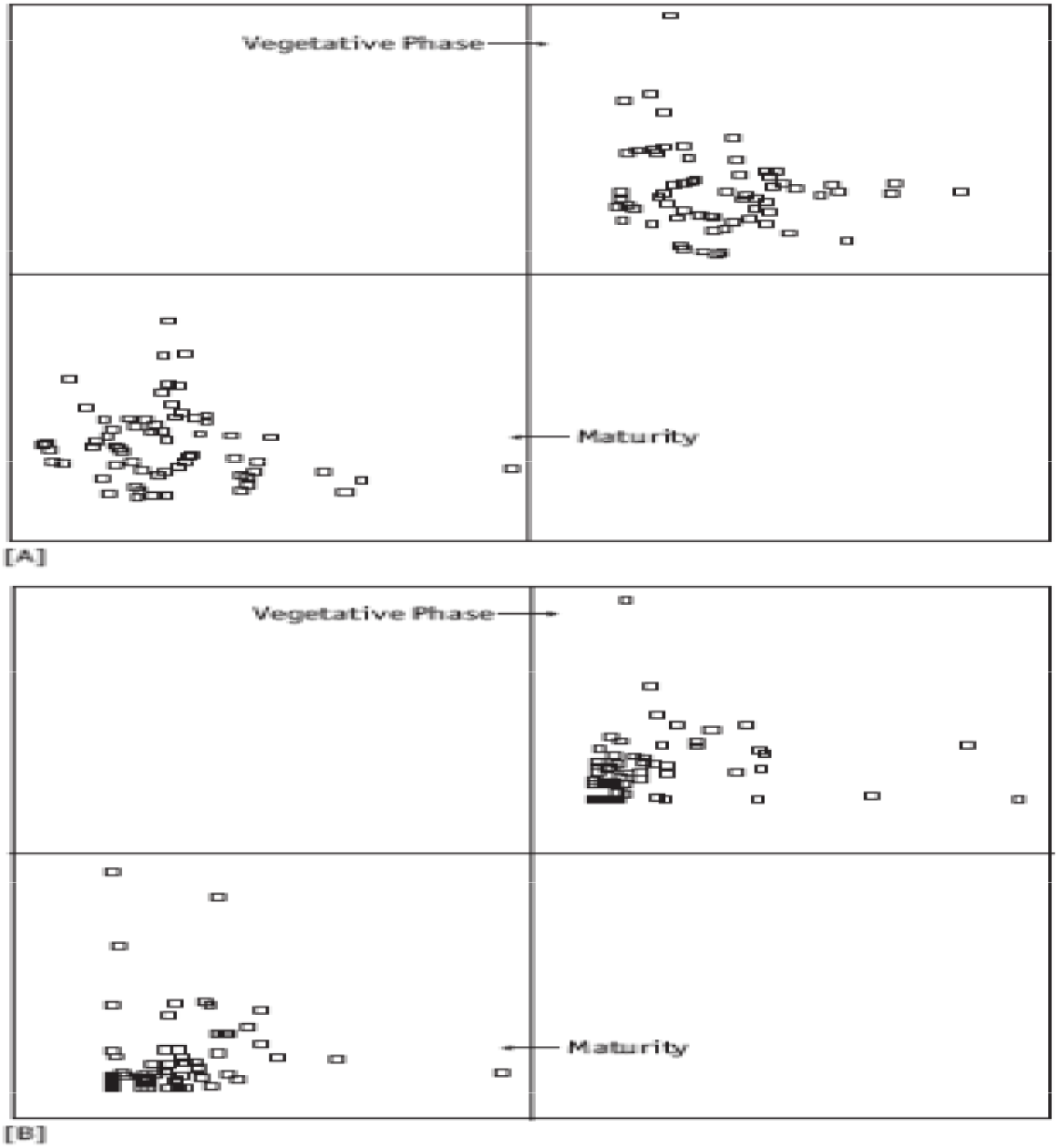
Scattered plot of withaferin-A content (% dry weight) showing distinctly high withaferin-A content in leaves (A) and roots (B) during vegetative phase of the plant.

**Table 3.** Range of Withaferin-A content (% dry weight) in *Withania somnifera* accessions.

|  | Minimum | Maximum | Mean | Std. Deviation |
| --- | --- | --- | --- | --- |
| 60 DoP Leaves | 0.2251 | 3.3821 | 1.0999 | ±0.5422 |
| 60 DoP Roots | 0.0003 | 0.7515 | 0.1243 | ±0.1230 |
| 150 DoP Leaves | 1.1657 | 1.2770 | 1.2138 | ±0.0572 |
| 180 DoP Leaves | 0.0056 | 1.9925 | 0.5943 | ±0.4030 |
| 180 DoP Roots | 0.0008 | 0.1489 | 0.0214 | ±0.0280 |

**Table 4.**
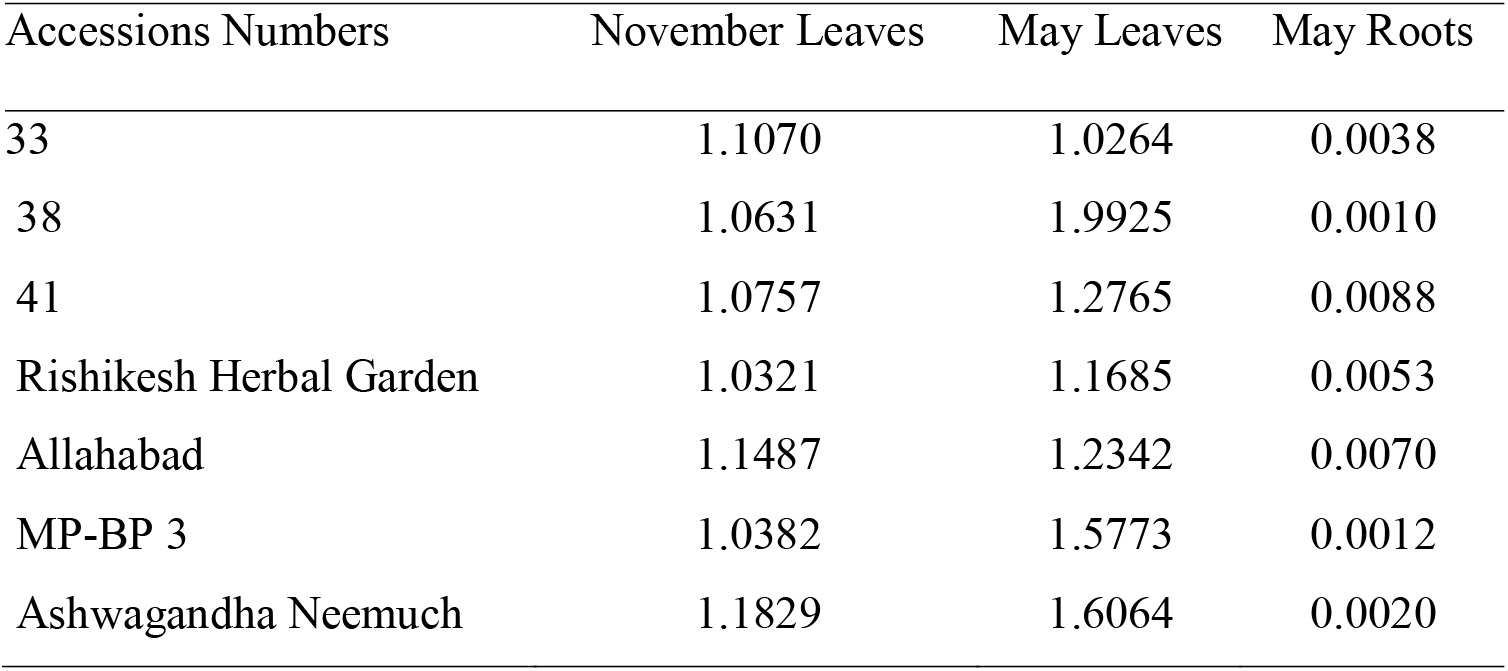
Accessions showing high Withaferin-A content (% dry weight)

| Accessions Numbers | November Leaves | May Leaves | May Roots |
| --- | --- | --- | --- |
| 33 | 1.1070 | 1.0264 | 0.0038 |
| 38 | 1.0631 | 1.9925 | 0.0010 |
| 41 | 1.0757 | 1.2765 | 0.0088 |
| Rishikesh Herbal Garden | 1.0321 | 1.1685 | 0.0053 |
| Allahabad | 1.1487 | 1.2342 | 0.0070 |
| MP-BP 3 | 1.0382 | 1.5773 | 0.0012 |
| Ashwagandha Neemuch | 1.1829 | 1.6064 | 0.0020 |

In Withania, plants with short height are also known as Nagori type and others having taller heights are known as Kashmiri type (Dr Sekhawat, Jodhpur University, personal communication). Origin of these nomenclatures is unknown. The Nagori type resembles the plant type of the cultivated varieties and the Kashmiri type resembles wild type. The plant type of the accessions growing at TERI’s field station broadly was of these two distinct types (Figure 7). ANOVA analysis revealed that the Withaferin-A content was significantly very high in the cultivated type than the wild type (Table 5).

**Figure 7.**
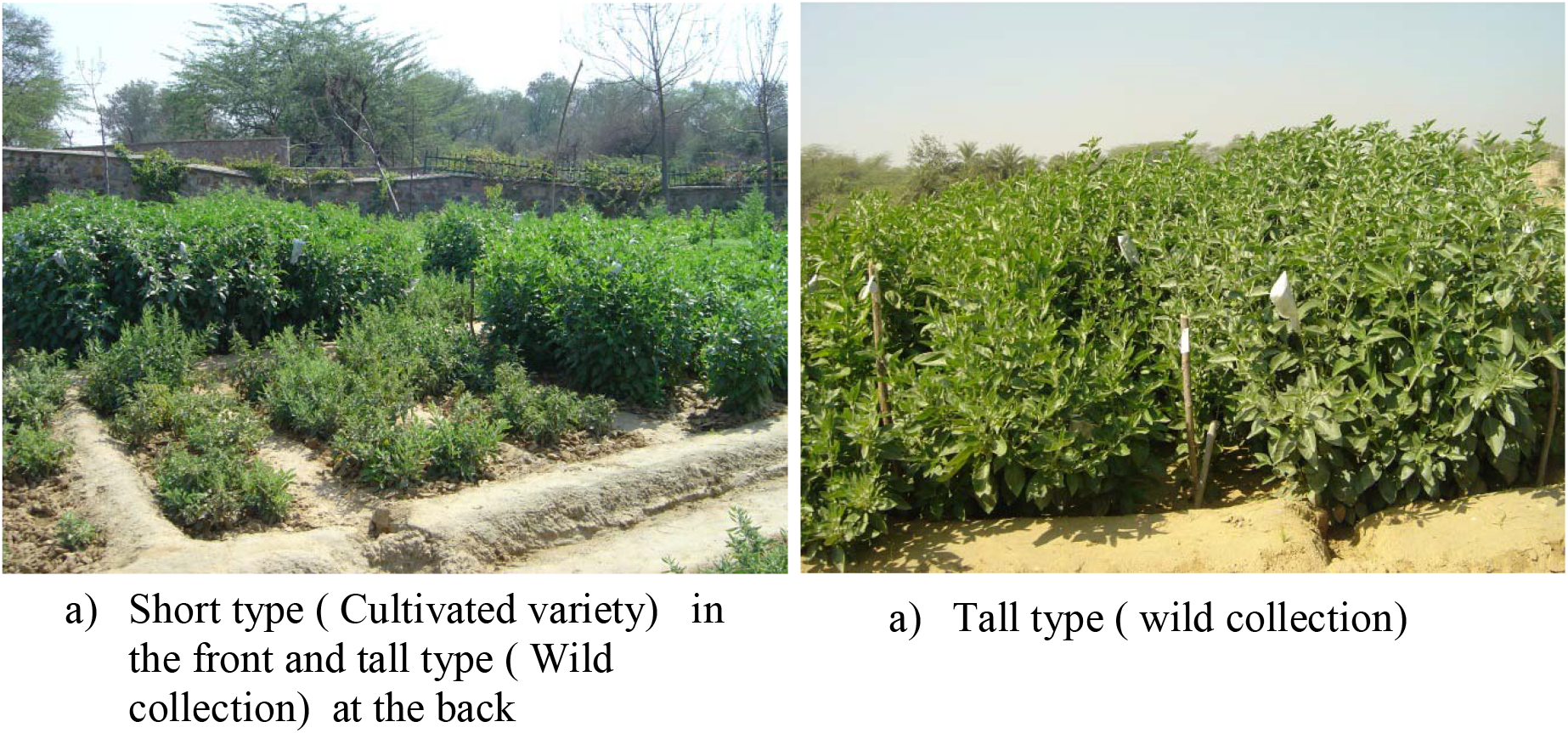
Two distinct morphotypes of Indian *Withania somnifera* germplasm.

**Table 5.** ANOVA showing effect of plant type-wild vs cultivated *Withania somnifera* on Withaferin-A content (% dry weight)

|  | Sum of Squares | df | Mean Square | F | Sig. |
| --- | --- | --- | --- | --- | --- |
| Between Groups | 4.314 | 1 | 4.314 | 18.661 | .000 |
| Within Groups | 14.795 | 64 | .231 |  |  |
| Total | 19.109 | 65 |  |  |  |

Difference in the Withaferin-A content in the wild type and the cultivated type were statistically highly significant. The population collected from different places differed significantly in Withaferin-A content (Table 6).

**Table 6.** ANOVA showing effect of location of collection of *Withania somnifera* on Withaferin-A content (% dry weight).

|  | Sum of Squares | df | Mean Square | F | Sig. |
| --- | --- | --- | --- | --- | --- |
| Between Groups | 5.817 | 9 | .646 | 2.723 | .010 |
| Within Groups | 13.293 | 56 | .237 |  |  |
| Total | 19.109 | 65 |  |  |  |

So far, limited efforts have been made to develop an analytical HPLC profile of the major constituents of *W. somnifera*. The HPLC methods reported earlier^7–11^ involved two to three step sample preparation and did not provide the leaf and root HPLC fingerprints. The HPLC method reported by Chaurasiya *et al*.^12^ involved tedious sample preparation and may not be suitable for routine analysis. The method reported in this paper was simple and useful not only for withanolide estimations but also for quality monitoring of the raw material used in drug preparation. Unlike Chaurasiya *et al*.^12^ who observed five major withanolides in leaf samples, we observed two major peaks in leaf fingerprint and many peaks in root fingerprint.

Withanolides have fairly wide geographic distribution and several chemotypes have been reported in literature based on the types and content of variously substituted steroidal lactones of withanolide series present in the species. The three chemotypes (I, II, III) reported so far in the Withania plants are reported from Israel^7,8^. Sicilian plants from Italy with absence of substitution at C-4, configurations 20S and 14βOH, and predominance of 17 α-OH compounds, belong to the Israelian chemotype III. The only difference is in the presence of withanolide-J as chief constituent as compared to the withanolide-E in Israelian species. Similar research studies on Sardinian plants led to the isolation of identical components as of Sicilian ^2^.

In India, *W. somnifera* grows throughout the drier parts and sub-tropical India. The plant is widely distributed in NorthWest part of India i.e. Bombay, Gujarat, Rajasthan, Madhya Pradesh, Uttar Pradesh, Punjab plains, which extends up to the mountain region of Punjab, Himachal Pradesh and Jammu.. Kirson et al. in 1969^13^ reported Withaferin A and withanone, as major constituents in the Indian *W. somnifera* plant collected from Delhi. Subramanian *et al*.^14^ (1971), isolated Withaferin A and a withanolide, C_28_H_38_O_6_, identical to the compound reported by Kirson, *et al*.^15^ (1970), from leaves of *W. somnifera* from Delhi region. After this, Chakraborti *et al*.^16^ (1974) reported that *W. somnifera* collected in West Bengal and Tamil Nadu differed chemotypically. The plant occurring in Tamil Nadu contained predominantly Withaferin A (I) where as in the West Bengal variety, 4β,20 α-dihydroxy-1-oxo-5 β, 6β epoxywith-2,24-enolide was the major constituent

Present study revealed the identification of pure Chemotype I in Indian germplasm which is not reported so far beside the intermediate of Chemotype I and Chemotype II. The study further revealed that the Withaferin content is highest during the vegetative growth of the plant, and leaves are the best source for extraction of Withaferin-A from withania plant. Further, it was also established that the Withaferin-A content in the leaves is not getting translocated to the roots at maturity of the plant as Withaferin-A content in roots also decreased at the time of maturity. Chaurasiya *et al*.^12^ have also reported that withanolides are synthesized in different parts of the plant (through the metabolic pathway) rather than being imported. Withaferin-A has emerged as an important withanolide having anticancer properties. Therefore, leaves during the vegetative phase of the plants can be a good source for harvesting this withanolide than roots. Plants yielding high Withaferin-A content can be developed as a new variety. Plants showing high production of Withaferin-A content irrespective of the age are also interesting and need further investigations.

## Conclusion

So far, limited efforts have been made to develop an analytical HPLC profile of the major constituents of *Withania somnifera*. The method reported in this paper was simple and useful not only for withanolide estimations but also for quality monitoring of the raw material used in drug preparation. Two major peaks were observed in leaf fingerprint and many peaks in root fingerprint. Present study revealed the identification of pure Chemotype I in Indian germplasm which was not reported so far beside the intermediate of Chemotype I and Chemotype II. The study further revealed that the Withaferin content was highest during the vegetative growth of the plant, and leaves were the best source for extraction of Withaferin-A from withania plant. Further, it was also established that the Withaferin-A content in the leaves did not translocate to the roots at maturity of the plant. This is because Withaferin-A content in roots decreased at that time. Leaves during the vegetative phase of the plants can serve as a good source for harvesting this withanolide rather than roots. Plants yielding high Withaferin-A contents can be developed as a new variety. Plants showing high production of Withaferin-A content irrespective of the age are also interesting and need further investigations.

## Acknowledgement

This work was supported by the Department of Biotechnology, Government of India.

## Notes

### Competing Interest Statement

The authors have declared no competing interest.

